# Distinct roles of PV and Sst interneurons in visually-induced gamma oscillations

**DOI:** 10.1101/2023.04.08.535291

**Authors:** Irene Onorato, Athanasia Tzanou, Marius Schneider, Cem Uran, Ana Broggini, Martin Vinck

**Affiliations:** Ernst Strüngmann Institute (ESI) for Neuroscience in Cooperation with Max Planck Society, 60528 Frankfurt, Germany; Max Planck Institute for Brain Research, 60438 Frankfurt, Germany; Donders Centre for Neuroscience, Department of Neuroinformatics, Radboud University Nijmegen, 6525 AJ Nijmegen, Netherlands

## Abstract

Sensory processing relies on interactions between excitatory and inhibitory neurons, which are often coordinated by 30-80Hz gamma oscillations. However, the specific contributions of distinct interneurons to gamma synchronization remain unclear. We performed high-density recordings from V1 in awake mice and used optogenetics to identify PV+ (Parvalbumin) and Sst+ (Somatostatin) interneurons. PV interneurons were highly phase-locked to visually-induced gamma oscillations. Sst cells were heterogeneous, with only a subset of narrow-waveform cells showing strong gamma phase-locking. Interestingly, PV interneurons consistently fired at an earlier phase in the gamma cycle (≈6ms or 60 degrees) than Sst interneurons. Consequently, PV and Sst activity showed differential temporal relations with excitatory cells. In particular, the 1st and 2nd spikes in burst events, which were strongly gamma phase-locked, shortly preceded PV and Sst activity, respectively. These findings indicate a primary role of PV interneurons in synchronizing excitatory cells and suggest that PV and Sst interneurons control the excitability of somatic and dendritic neural compartments with precise time delays coordinated by gamma oscillations.

## Introduction

The cortical microcircuit has a canonical composition of excitatory neurons and various types of GABAergic interneurons. E/I (excitatory-inhibitory) interactions are essential to sensory processing (Adesnik et al., 2012; Isaacson and Scanziani, 2011; Gentet et al., 2012; Muñoz et al., 2017; Yu et al., 2019) and give rise to rhythmic network activity in various frequency bands (Buzsáki and Draguhn, 2004; Wang, 2010). In the visual cortex, neurons often exhibit synchronized firing at a fast time scale orchestrated by the gamma rhythm (Gray et al., 1989). This rhythm is thought to be involved in various processes including attention, predictive processing, phase coding and assembly formation (Singer and Gray, 1995; Singer, 1999; Fries, 2005; Fries et al., 2007; Vinck et al., 2013; Pesaran et al., 2002; Uran et al., 2022; Peter et al., 2021; Speed et al., 2019; de Almeida et al., 2009), and also serves as a marker for neurodegenerative and psychiatric diseases associated with dysfunctional E/I interactions (Uhlhaas and Singer, 2010; Ray, 2022). Evidence suggests that the generation of gamma rhythms depends on E/I interactions, yet the distinct roles of different interneuronal classes in the generation of cortical gamma oscillations are still not fully understood (Tukker et al., 2007; Hahn et al., 2022; Börgers et al., 2008; Veit et al., 2017; Chen et al., 2017; Hakim et al., 2018; Cardin et al., 2009; Cardin, 2018; Kopell et al., 2000).

The main sources of inhibitory inputs onto cortical excitatory neurons are Parvalbumin (PV) and Somatostatin-expressing (Sst) interneurons, which mostly correspond to the fast-spiking basket cells and Martinotti cell-types, respectively. PV interneurons preferentially target the perisomatic compartment of pyramidal neurons, while Sst interneurons preferentially target their dendrites (Rudy et al., 2011a; Pfeffer et al., 2013; Cardin, 2018). Hence, PV interneurons gate excitatory synaptic inputs globally while Sst interneurons gate specific dendritic segments (Chiu et al., 2013; Cardin, 2018). In classic models of cortical gamma, the synchronization of excitatory neurons results from rhythmic inhibition mediated by PV cells (Buzsáki and Wang, 2012; Börgers et al., 2005; Vinck et al., 2013; Csicsvari et al., 2003; Cardin et al., 2009; Perrenoud et al., 2016). Such a role would fit with the observation that PV interneurons are highly responsive to transient increases in excitatory inputs due to their rapid dynamics and depressive E-to-PV synapses (Jouhanneau et al., 2018; Cardin, 2018). Computational models further predict that tonic inhibition from other GABAergic sub-types, e.g. Sst neurons, might facilitate the emergence of gamma by reducing the excitability of PV and excitatory cells (Börgers et al., 2008).

Recent studies however suggest an alternative model for visually-induced gamma oscillations, in which neuronal rhythmic activity results from reciprocal interactions between Sst and excitatory cells (Veit et al., 2017; Hakim et al., 2018). This model builds on the observation that visually-induced gamma oscillations typically emerge for large, homogeneous stimuli (Vinck and Bosman, 2016; Veit et al., 2017; Uran et al., 2022) which tend to activate Sst interneurons while suppressing PV interneurons (Adesnik et al., 2012; Keller et al., 2020) (but see Dipoppa et al. (2018)). Interestingly, recent work also suggests an important role of burst firing of excitatory cells in the generation of gamma (Onorato et al., 2020), which would fit with the observation that Sst interneurons are effectively driven by bursts but also control burst firing (Gentet et al., 2012; Royer et al., 2012; Murayama et al., 2009; Cardin, 2018; Jouhanneau et al., 2018). Hence, gamma-rhythmic activity may represent the timed control of burst firing via Sst interneurons.

At present, evidence from causal experiments is compatible with both models as photo-suppression of either Sst and PV interneurons can lead to strong reductions in V1 gamma-amplitude (Veit et al., 2017; Hakim et al., 2018; Chen et al., 2017). Hence, data suggests both Sst and PV interneurons contribute to gamma oscillations, but it remains to be determined what these contributions precisely are. A fundamental understanding of the generation of gamma and the distinct contributions of Sst and PV interneurons requires (1) measuring the phase-locking strength and the timing at which these specific interneurons are recruited in the gamma cycle; and (2) recording excitatory neurons simultaneously. To this end, we performed high-density recordings from area V1 in awake mice and used transgenic mice lines to render PV and Sst interneurons light-sensitive, which allowed us to selectively tag them in awake mice V1. We then analyzed the spike-timing and phase-locking of various groups of interneurons, and examined how their activity is coordinated with that of excitatory neurons within the gamma cycle.

## Results

### Identification of PV and Sst interneurons using optogenetics

We recorded LFPs and spiking activity from area V1 in mice using high-density laminar probes (see Methods; Fig. 1a). Mice were head-fixed while placed on a running wheel and passively viewed full-field drifting-grating stimuli. To record from specific GABAergic interneuron sub-types, we crossed the PV and Sst Cre-lines to the Ai32 line, yielding expression of Channelrhodopsin-2 (ChR2) in either PV or Sst interneurons (Fig. 1a). The cortical tissue was illuminated with square-wave (i.e. DC) light stimuli of 1 s duration. Optotagged (i.e. ChR2expressing) neurons were identified by their increased firing rates during the presentation of the light stimulus (see methods, Fig. 1b-c). In the same sessions, we also recorded from (putative) excitatory neurons that were identified by their broad waveforms (n = 2156, Fig. S1a).

**Figure 1:**
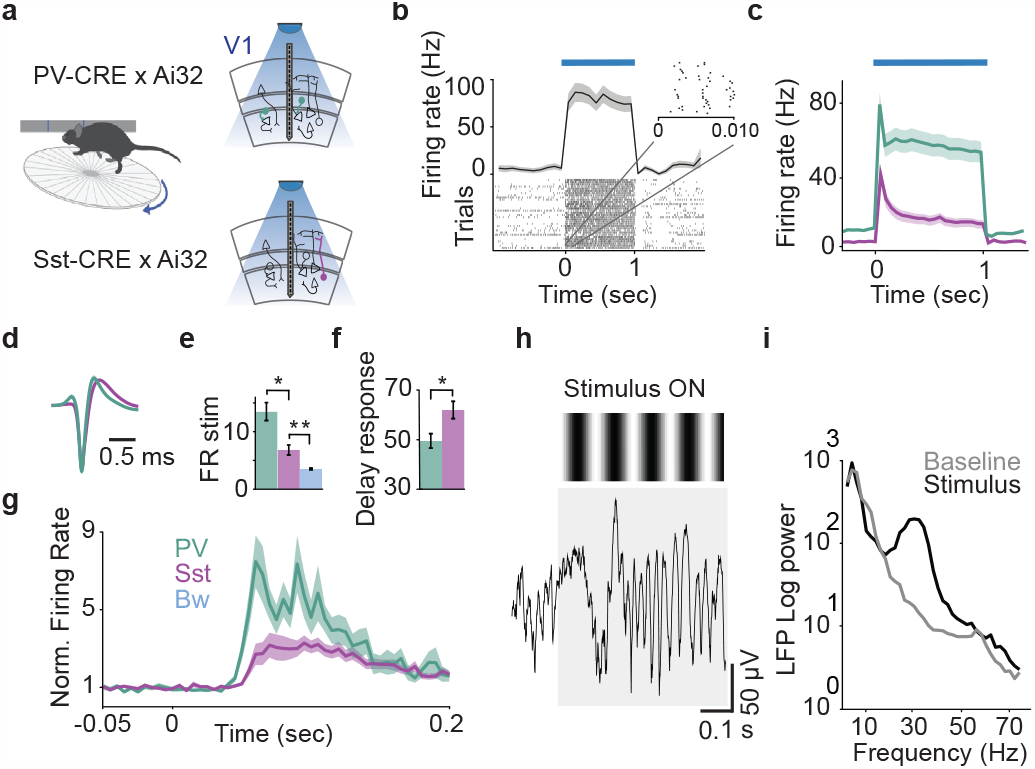
Recording paradigm and identification of PV and Sst interneurons. **a** Recordings of LFPs and spikes from area V1 were made with high-density silicon probes (Neuropixels and Cambridge Neurotech). PV or Sst interneurons were identified using transgenic lines (Ai32 ×PV-CRE and Ai32 ×SOM-CRE). Shown is an example response of a neuron to light onset. Mice were placed on a running wheel, while we presented drifting-grating stimuli. **b** Example of an opto-tagged neuron with increased firing rates during 1 s of light stimulation. **c** Average light responses of PV (green, n = 73) and Sst interneurons (magenta, n = 92). **d** Average normalized spike waveforms of PV and Sst interneurons. **e** Average firing rates during the initial stimulus transient. PV interneurons had a higher firing rate than Sst interneurons (p = 0.0001, two-sample t-test), which had higher rates than broad-waveform, putative excitatory neurons (Sst vs. Bw, p = 1.5 ×10^−7^). **f** Comparison of average stimulus latency response (see Methods) between Pv and Sst interneurons (PV vs. Sst: 49 vs. 60 ms, p = 0.0066; two-sample t-test). **g** The average peri-stimulus-time-histogram, which was normalized by dividing by the baseline firing rate. **h** Example of raw LFP trace around visual stimulus onset. **i** Example LFP power spectrum during drifting gratings presentation (black line) and the gray-screen baseline (gray line). **c-i** Error bars and shaded lines indicate s.e.m.

The identified PV and Sst interneurons showed comparable laminar distributions (Fig. S1b). PV and Sst interneurons typically had narrow(Nw) and broad-waveform (Bw) action potentials, respectively (Fig. 1d and Fig. S1c). Both PV and Sst neurons showed higher discharge rates than Bw excitatory cells, both during the stimulus (Fig. 1e) and baseline period (Fig. S1e. Furthermore, PV interneurons had higher baseline (i.e. pre-stimulus) firing rates than Sst interneurons (Fig. S1e).

Both Sst and PV interneurons had enhanced firing rates during the stimulus presentation (Fig. 1g). PV interneurons were more visually driven (Fig. S1f), and their evoked responses had shorter onset latencies than Sst interneurons (11 ms time difference, Fig. 1f). The observed differences between PV and Sst neurons in terms of spike waveforms and firing properties are broadly consistent with previous studies (Miri et al., 2018; El-Boustani and Sur, 2014; Muñoz et al., 2014; Kim et al., 2016; Jang et al., 2020; Senzai et al., 2019; Perrenoud et al., 2016; Yu et al., 2019; Gentet et al., 2012; Senzai et al., 2019; Estebanez et al., 2017).

### Comparison of phase-locking properties between PV and Sst interneurons

During the presentation of drifting-grating stimuli, we observed prominent gamma oscillations in the LFP, as previously reported by Veit et al. (2017), (Fig. 1h). These gamma oscillations were characterized by a peak in the LFP’s power spectral density around 30 Hz, with a clear elevation in stimulus-induced power as compared to baseline (Example session: Fig. 1i; population-average: Fig. S1d). Note that for LFP analyses, we discarded trials in which the network showed strong 4-8 Hz oscillations (Gao et al., 2021; Speed et al., 2019; Senzai et al., 2019), as these spindle-like events disrupted gamma-synchronization.

To determine the contribution of different cell classes to the generation of gamma oscillations, we used the classic approach of determining the phase-locking of spikes to the local LFP gamma oscillations (Csicsvari et al., 2003; Vinck et al., 2013; Tukker et al., 2007; Onorato et al., 2020; Klausberger et al., 2003; Bragin et al., 1995).

Neurons showed consistent spike phase-locking to LFP gamma oscillations, which is illustrated in two representative PV and Sst example neurons. In the example of the PV interneuron, spikes tended to consistently occur at a specific phase of the gamma cycle (Fig. 2a). These phases were quantified with the wavelet transform of the V1 LFP signal centered on each spike. The consistent clustering of spike-LFP phases resulted in clear side-lobes in the spike-triggered LFP average (STA, Fig. 2a). To determine the strength of spike phase-locking to the LFP, we computed the PPC measure (pairwise phase consistency). The PPC measure is proportional to the squared resultant length, but is unbiased by spike count and not affected by firing-rate history effects like bursting (Vinck et al., 2012). The PPC spectrum of the example PV interneuron showed a narrow-band peak around 30 Hz (Fig. 2a). By contrast, the Sst example neuron fired spikes in a broader range of gamma phases, which resulted in an STA without prominent oscillatory side-lobes and lower PPC values at gamma frequencies (Fig. 2b).

**Figure 2:**
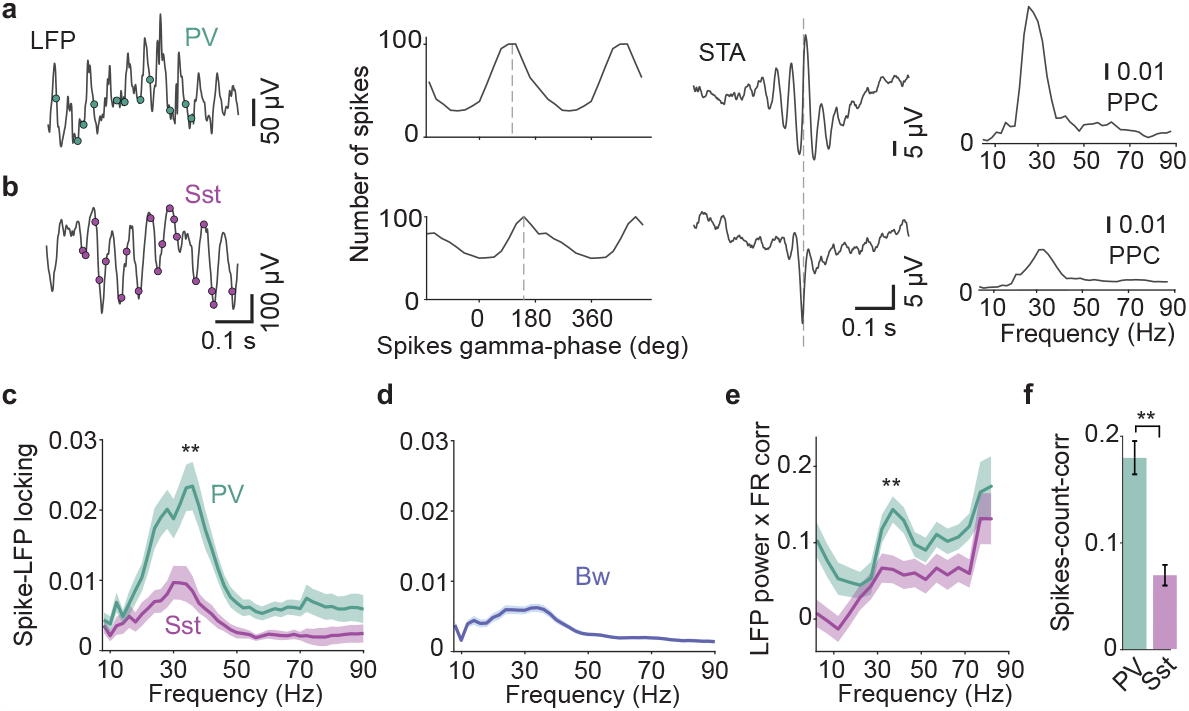
PV and Sst interneurons show differences in gamma phase-locking and firing correlations to network activity. **a**) Example PV interneuron. Leftto-right: LFP trace with spikes of PV interneuron (green dots); histogram of the spike gamma-phase distribution; spike-triggered-average of the LFP; spike-LFP phase-locking quantified by the pairwise phase consistency (PPC) measure. The PPC is unbiased by spike count and not affected by spike train history effects like bursting. **b**) Same as **a**, but for a representative Sst interneuron. **c**) Average PPC for PV and Sst interneurons. PV interneurons have higher PPC values in the gamma-frequency range than Sst interneurons (mean difference in 25-45 Hz range: p = 2 ×10^−4^, two-sample t-test). **d**) Bw (putative excitatory) neurons have lower PPC values than PV (p = 0.008) but not Sst cells (p = 0.25). **e**) Mean correlation value between a neuron’s firing rate and spectral LFP power. PV interneurons show strong correlations in the gamma-frequency range than Sst interneurons peaking between 37 and 44 Hz (p = 0.003; two-sample t-test). **f**) Noise correlation between a neuron’s spike count and the spike counts of simultaneously recorded neurons. The correlations were computed across trials, separately for PV and Sst interneurons. PV interneurons have stronger correlations than Sst interneurons (p = 1.6 ×10^−7^; two-sample t-test). **c-f** Error bars and shaded lines indicate s.e.m.

To investigate differences in phase locking at the population level, we computed the average spike-LFP phase-locking across neurons, separately for the Sst and PV interneuron populations. Both PV and Sst interneurons showed a peak in the spikeLFP phase-locking spectrum around 30Hz, indicating that neural firing was synchronized with the local LFP gamma oscillation (Fig. 2c). PV interneurons showed significantly stronger gamma phase-locking strength than both Sst interneurons (2.5 fold increase) and Bw excitatory neurons (3.2 fold; Fig. 2c-d). By contrast, we did not find a difference in phase-locking strength between Sst interneurons and Bw excitatory neurons (Fig. 2c-d). Stronger phase-locking in PV than Sst neurons was also found when restricting the analysis to neurons in superficial layers, and when using different criteria for opto-tagging (Fig. S2). Furthermore we replicated the observed difference in phase-locking between PV and Sst interneurons in a smaller dataset from the Allen Brain Institute (Fig. S2).

To further investigate the contributions of PV and Sst interneurons to the local gamma rhythm, we investigated the relationship between their instantaneous firing rates and the gamma-amplitude of the LFP. To this end, we computed the Spearman correlation between the trial-by-trial change in spike rates with the LFP power at different frequencies (Fig. 2e). Both PV and Sst neurons showed positive rate gamma oscillations, which is illustrated LFP-power correlations in the gamma-frequency range. However, only PV interneurons showed a gamma peak in the rate × LFP-power correlation spectrum, which was absent in Sst interneurons (Fig. 2e). We note that this result differs from a previous study, in which a smaller sample of Sst neurons showed a stronger positive correlation with gamma LFP amplitude than the firing rates of PV neurons (Veit et al., 2017).

The above findings showed that, compared to Sst interneurons, PV interneurons have more precise phase-locking and their firing rates are more positively correlated to the amplitude of the gamma network oscillation. We therefore wondered if the firing rates of PV interneurons are also more correlated with the firing rates of other neurons in the network on longer time-scales. To investigate this, we computed the spike-count noise correlation between each opto-tagged interneuron and all the other neurons that were recorded from the same laminar probe. We found that the firing rates of PV interneurons were more (positively) correlated with the firing rates of the other recorded neurons than was the case for Sst interneurons (fig 2f).

Together, these findings show that PV interneurons rather than Sst interneurons exhibit the strongest temporal coordination local network activity, both at the time-scale of a gamma cycle and at longer time scales. These findings suggest that PV interneurons are the main source of gamma-rhythmic inhibitory inputs onto excitatory neurons.

### Relation between firing patterns of excitatory neurons and GABAergic interneurons

Next, we asked how the spiking activity of PV and Sst interneurons is coordinated with Bw excitatory neurons. Classic models of cortical gamma at lower frequencies are based on the PING model (Hasenstaub et al., 2005; Wang, 2010; Tiesinga and Sejnowski, 2009; Kopell et al., 2000; Wilson and Cowan, 1972; Buzsáki et al., 2012; Atallah and Scanziani, 2009; Cardin et al., 2009; Csicsvari et al., 2003). In the PING model, each gamma cycle starts with an increase in the firing of excitatory neurons, which then leads to an increase in the firing of inhibitory interneurons, causing a subsequent decrease in the firing activity of both excitatory neurons and inhibitory interneurons.

To gain further insight into the interactions of PV and Sst interneurons with the local network, we characterized the distribution of spike-LFP phases across neurons (Csicsvari et al., 2003; Vinck et al., 2013). For each neuron separately, we computed its preferred (i.e. average) phase-of-firing in the gamma cycle. We then constructed a histogram of preferred spike gamma-phases, separately for the PV and Sst populations (Fig. 3a). PV cells tended to fire relatively early in the gamma cycle, with their spikes falling on the gamma cycle’s descendent phase (≈130 degrees, i.e. 50 degrees before the trough). Compared to PV interneurons, the preferred gamma-phase of Sst interneurons showed a substantial phase-delay of about 60 degrees, with spikes falling shortly after the gamma cycle’s trough (Fig. 3a; mean phase difference PV and Sst).

**Figure 3:**
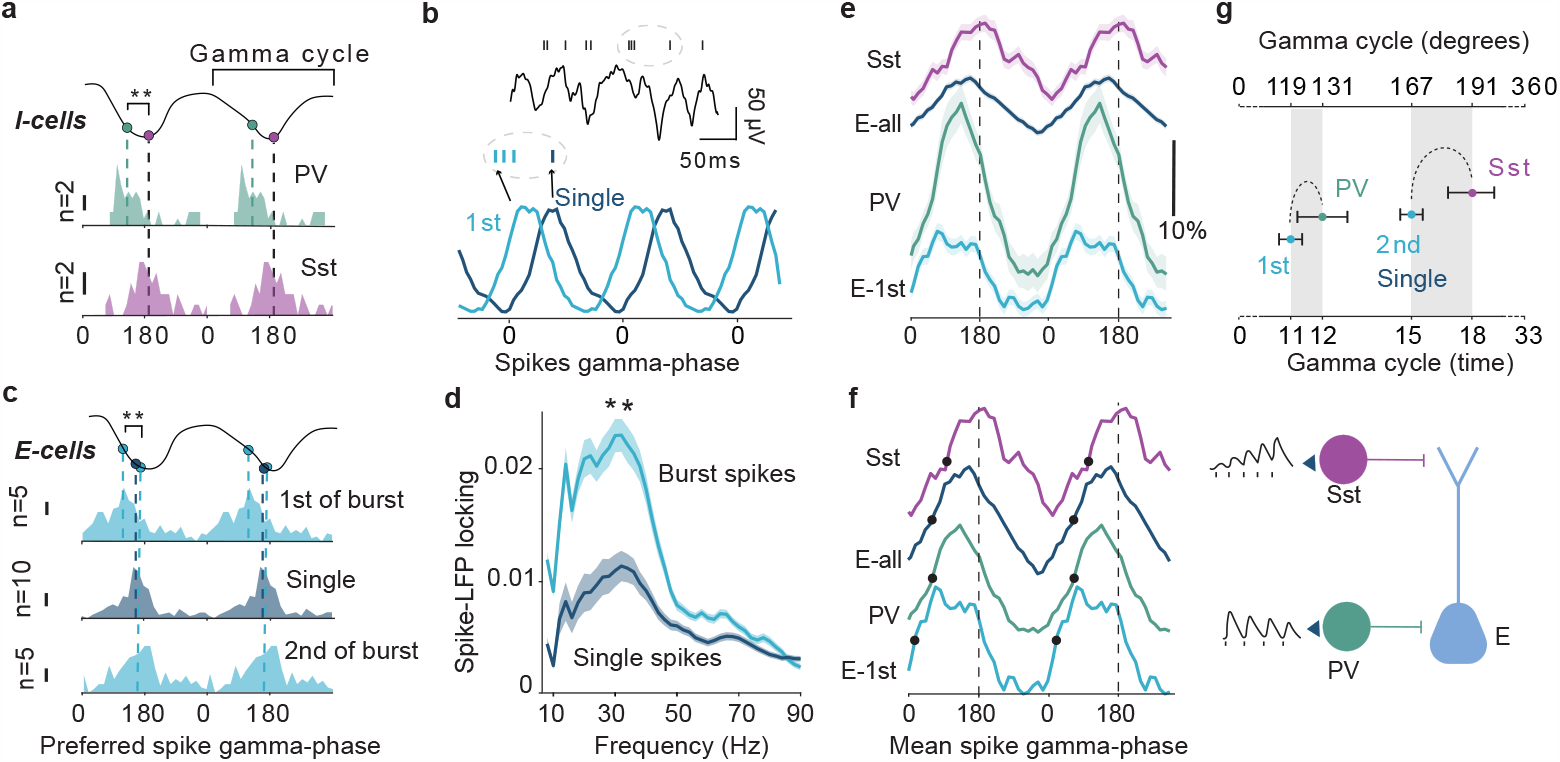
Delayed firing of SSt interneurons in the gamma cycle compared to PV interneurons. **a**) Comparison of gamma-phase distributions across neurons between PV and Sst interneurons. For each neuron, we computed its preferred (i.e. average) gamma-phase. Shown is the histogram across neurons for PV (n = 55) top and Sst (n = 42). PV interneurons fired significantly earlier in the gamma cycle (131 degrees) than Sst cells (190 degrees; p < 0.005; permutation test). **b**) Example LFP trace and the spiking activity of a Bw neuron. Spikes are divided into burst spikes (n = 770, inter-spike-interval <6ms) and single spikes (NBS,n = 750, inter-spike-interval >10ms). Histogram of the gamma-phase distribution for burst and single spikes. **c**) Distribution of the preferred spike gamma-phase for the first (top plot) and second (bottom plot) spikes of burst events, and the single spikes (middle plot). The 1st spikes of the burst occurred earlier in the gamma cycle than single spikes (p < 0.0005, permutation test) and the 2nd spike of the burst (p < 0.001). There was no difference in phase between single spikes and the 2nd spike of burst (p > 0.05; permutation test). **d**) Spikes-LFP phase locking (PPC) for the burst spikes (average PPC values between first and second burst spikes) and single spikes. Burst spikes had stronger phase locking (2.1 folds) in the gamma-frequency range (25-40 Hz, p = 7.5 gamma oscillations, which is illustrated 10^−6^; two-sample t-test). **e**) Comparison of average phase-density of spiking activity between neural populations. Phase histograms were first computed for each neuron separately, normalized to the mean, and then averaged across cells. **f**) Left: shown are the curves from **e** normalized to the maximum. Filled circles correspond to time points where cells reach 50% of the maximum. Right: illustration of depressing (E-to-PV) and facilitating (E-to-Sst) synaptic properties, and the projections of PV and Sst interneurons to excitatory cells. **g**) Summary of preferred spike phase for PV, Sst, 1st spike of the burst and single spikes. **d-g** Error bars and shaded lines indicate s.e.m.

Similar results, i.e. a phase advance of PV relative to Sst interneurons, were obtained in an independently acquired dataset from the Allen Brain observatory, in which a smaller sample of PV and Sst interneurons was identified with a similar methodology. Furthermore, the phase advance of PV interneurons was also observed when restricting to superficial units (Fig. S2).

Interestingly, the average preferred gamma-phase of excitatory neurons (at 162 degrees) was slightly delayed relative to the PV interneurons. A phase advance of PV interneurons relative to excitatory neurons seems inconsistent with the PING model. To further understand the implications of this phase advance, we performed two kinds of analyses.

First, we considered that the activity of excitatory neurons can be highly heterogeneous (Onorato et al., 2020; Bartho et al., 2004; Vinck et al., 2016). In particular, the spikes of excitatory neurons can be divided into burst (defined as consecutive spikes within 6 ms) and single (i.e. non-burst) spikes (more than 10 ms between the previous and next spike) (Fig. 3b). Burst spikes comprised 7.3% of all the spikes fired by Bw cells, and were substantially more phase-locked than single spikes (Fig. 3d). We found that the first spike of the burst showed, on average, a significant phase advance relative to the PV interneurons of about 1 ms (Fig. 3e). In addition, the first spike of the burst was also phase-advanced relative to single spikes and the subsequent spikes in the burst (Fig. 3c). Thus, the first spike of the burst may be an important event to ignite the gamma cycle, because of its strong phase-locking and early phase preceding the PV interneurons (Fig. 3e and Fig. 3f). By contrast, the spiking of Sst interneurons followed the single and 2nd spike of the burst (Fig. 3e and Fig. 3f). Hence, Sst neurons may be driven by a broader set of excitatory inputs that include the second spike of burst events. In fact, the average phase of the second spike of the excitatory-neuron bursts preceded the Sst neurons by about 2-3 ms (Fig. 3e). Because Sst neurons have facilitatory synapses, they are particularly sensitive to repeated stimulation (e.g. bursts) (Cardin, 2018; Jouhanneau et al., 2018), which offers a potential explanation for their delayed firing in the gamma cycle.

Second, we considered that the histogram of preferred phases across neurons does not necessarily reveal the precise rise of inhibition and excitation within the gamma cycle. We therefore performed a complementary analysis to examine the distribution of phases, by first constructing a normalized (to the mean) gamma-phase histogram per neuron, and then averaging these histograms across neurons. PV interneurons showed a larger phase modulation-depth as compared to Sst neurons and excitatory neurons (see Fig. 2), which is consistent with the higher PPC values in PV interneurons. Moreover, this analysis replicated the clear gamma-phase advance of PV neurons as compared to Sst neurons, and the advance of the first spike of the burst relative to PV interneurons. Importantly, when considering all spikes fired by excitatory neurons together, it can be seen that the phase distributions of excitatory neurons and PV interneurons have different shapes. Specifically, the initial rise in spiking in excitatory cells coincides precisely with the rise in PV interneuron activity. However, PV interneurons show a relatively steep decline after reaching a peak, while the firing of excitatory neurons is prolonged in the gamma cycle (Fig. 3e-f). Thus, considering the heterogeneity of spikes fired by excitatory neurons and their behavior at the onset of the gamma cycle, the data is overall compatible with a PING mechanism between excitatory neurons and PV interneurons. Our findings suggest that PV interneurons are the main source of rhythmic inhibition gating excitatory inputs from all dendritic compartments. However, the delayed firing of Sst interneurons suggests a distinct contribution to the temporal control of excitatory cells. In particular, our analyses indicate that in each gamma cycle, perisomatic inhibition is followed by delayed dendritic inhibition mediated by Sst interneurons. Similar to what was observed after the onset of visual stimuli (Fig. 1), Sst interneurons may be recruited more slowly and especially by burst events due to facilitatory E-to-Sst synapses. The delayed dendritic inhibition roughly coincides with the timing of burst spikes and may therefore play an important role in controlling burst-spikes (Gentet et al., 2012; Royer et al., 2012; Murayama et al., 2009) (see Discussion).

### Sst subtypes

Sst interneurons are a heterogeneous cell class that includes sub-types with different morphological and physiological features (Muñoz et al., 2014, 2017; Nigro et al., 2018; Naka et al., 2019; Kim et al., 2016; Ma, 2006; Kvitsiani et al., 2013; McGarry, 2010). We therefore wondered if the Sst population contained distinguishable sub-types. To investigate this, we examined the action-potential waveform distribution of the Sst neurons. We found that the Sst interneurons included both Nw and Bw cells, consistent with previous reports (Muñoz et al., 2014; Kim et al., 2016; Nigro et al., 2018; Kvitsiani et al., 2013), (Fig. 4b). Bw Sst interneurons (≈ 71%) constituted the largest group, and the two types of Sst neurons showed comparable laminar distributions (Fig. S3a). We then repeated the main analyses reported above for the Nw and Bw Sst interneurons separately. Firing rate responses of Nw and Bw Sst interneurons to the visual stimulus had similar latencies (Fig. S3b-c). Nw Sst interneurons had higher firing rates, both during the stimulus presentation period and the baseline period (Fig. 4a and fig S3d), in agreement with previous studies (Kim et al., 2016).

**Figure 4:**
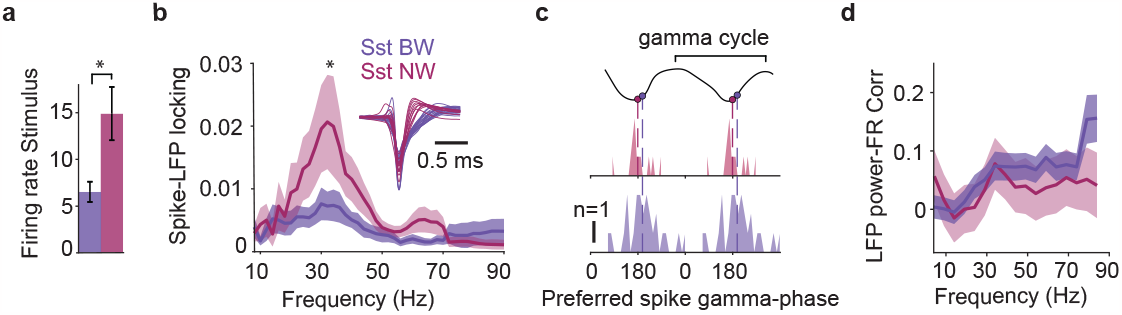
Narrow and broad waveform Sst show distinct functional properties. **a**) Average firing rate during the late stimulus period (time window between 0.4 and 1.8 seconds from the stimulus onset). Nw (n = 21) have higher firing rates than Bw (n = 54) Sst interneurons (p = 0.0012, two-sample t-test). **b**) Average spikes-LFP phase locking (PPC). Nw Sst have higher gamma PPC values than Bw Sst cells (at 25-40 Hz: p = 0.037; two-sample t-test). Inset shows Sst waveforms for Bw and Nw groups. **c**) Distribution of preferred gamma phases across neurons. There is no significant difference between Nw Sst (n = 28) and Bw Sst (n = 17) cells (p = > 0.05; permutation test). **d**) Mean correlation value between a neuron’s firing rates and LFP spectral power. There is no significant difference in correlation values at gamma frequencies (at 37-42 Hz, p = 0.64; two-sample t-test). **a-d** Error bars and shaded lines indicate s.e.m.

To investigate whether Nw and Bw Sst subtypes differed in their phase-locking properties, we computed the spike-LFP PPC value for each Sst neuron as above. We observed substantially stronger phase-locking in Nw Sst interneurons as compared to Bw Sst interneurons (Fig. 4b). However, both Sst subtypes showed similar spike-phase distributions in the gamma cycle (Fig. 4c). We further computed the correlation of Bw and Nw Sst firing rates with the LFP gamma-amplitude. This analysis did not show differences between the two groups of Sst interneurons (Fig. 4d).

Thus, Nw Sst interneurons exhibit functional similarities to PV interneurons, whereas they differ from Bw Sst interneurons in terms of their high firing rates, narrow-waveform action potentials, and strong synchronization to the gamma phase. However, the Nw Sst interneurons exhibited similar behavior to Bw Sst interneurons and were distinct from PV interneurons, in the sense of delayed firing in relation to the stimulus onset and within the gamma-cycle.

## Discussion

Sensory processing relies on E/I interactions, which are coordinated at the time-scale of gamma oscillations (Buzsáki and Wang, 2012; Tiesinga and Sejnowski, 2009; Bartos et al., 2007; Hasenstaub et al., 2005; Csicsvari et al., 1999). To determine how different GABAergic sub-types interact with excitatory neurons during gamma oscillations, we recorded from optogenetically identified PV and Sst interneurons in area V1 of awake mice. PV and Sst interneurons are the main sources of perisomatic and dendritic inhibition of excitatory neurons, respectively (Rudy et al., 2011a). Our first main finding is that PV interneurons were substantially (several fold) more phaselocked to gamma oscillations than excitatory neurons and Sst interneurons. Our second main finding, is that PV interneurons consistently fired at an earlier gamma phase (about 6ms or 60 degrees) than Sst interneurons. To gain insight into the circuit function, we further analyzed the phase delay in the activation of PV and Sst interneurons in comparison with the Bw cells. The firing of PV interneurons in the gamma cycle coincided with the average rise in spikes fired by excitatory neurons. But, the onset spikes of bursts fired by excitatory neurons, which were strongly gamma phase-locked, showed a clear phase advance over PV interneurons. By contrast, the firing of Sst interneurons typically followed the 2nd spike of excitatory neurons’ bursts. Finally, we identified a smaller subset of Sst interneurons with narrow-waveform action potentials. These neurons were strongly gamma phase-locked, but fired at a late gamma phase, similar to broad-waveform (Bw) Sst neurons.

In sum, these findings indicate that PV interneurons and a subclass of narrow-waveform Sst interneurons are the main sources of gamma-rhythmic inhibition of excitatory neurons. Perisomatic inhibition mediated by PV interneurons occurs at an early phase in the gamma cycle, whereas dendritic inhibition mediated by Sst interneurons arrives at a later gamma phase and is temporally coordinated with the burst spikes of excitatory neurons. The time delay between the activation of Sst interneurons and burst spikes in the gamma cycle is compatible with a bidirectional interaction: Sst interneurons inhibit bursts in excitatory neurons (Royer et al., 2012; Gentet et al., 2012), but are also effectively driven by burst spikes (Jouhanneau et al., 2018). Thus, PV and Sst interneuron control the excitability of somatic and dendritic neural compartments with precise time delays coordinated by gamma oscillations.

### Distinct roles of PV and Sst interneurons

Based on their strong gamma phase-locking, we conclude that PV interneurons are the main source of rhythmic inhibition that controls the spike timing of excitatory neurons. Such a role should be facilitated by two further properties of PV interneurons: (1) PV interneurons make highly divergent projections to a large number of excitatory neurons (Salkoff et al., 2015; Buzsáki and Wang, 2012); (2) Peri-somatic inhibition provides powerful control over neural firing, by hyperpolarizing the soma directly and shunting excitatory inputs from all dendritic compartments (Vu and Krasne, 1992; Koch, 2004).

The visually-induced gamma rhythm that we observed likely originates from a PING model, as the cycle duration is relatively long (about 33-40 ms). By contrast, the ING (Interneuron Network Gamma) model predicts gamma oscillations at higher frequencies dictated by the time constants of GABAa currents (Wang, 2010). In PING models, it is typically assumed that excitatory neurons exhibit a phase advance over inhibitory neurons, which has been observed in several studies (Hasenstaub et al., 2005; Vinck et al., 2013; Csicsvari et al., 2003; Börgers and Kopell, 2003; Salkoff et al., 2015). Surprisingly, we found that PV interneurons fired very early in the gamma cycle, preceding the average preferred phase of excitatory neurons. This seems, *prima facie*, difficult to reconcile with the PING model, and to suggest an ING model. However, further analyses indicated that in the gamma cycle, the initial rise in PV-interneuron activity is very closely aligned with the rise in activity of excitatory cells. Furthermore, our data indicate that burst events may be important in igniting the gamma cycle, as burst spikes showed strong phase-locking, and the 1st spike of the burst clearly preceded the PV interneurons in time. Finally, it needs to be considered that E-to-PV connections are very rapid (Jouhanneau et al., 2018).

In contrast to PV interneurons, Sst interneurons were relatively weakly phase-locked to gamma LFP oscillations, and their spikes were fired at a late gamma phase. The relatively weak gamma phase-locking of Sst interneurons fits well with the observation that compared to PV interneurons, these neurons respond with less temporal precision to synaptic inputs and have longer membrane time constants (Jouhanneau et al., 2018; Cardin, 2018). The observation of strong PV gamma-locking and weak Sst gamma-locking suggests two roles of Sst neurons in visually-induced gamma-band activity:

(1) *Borgers*/*Kopell model*. In the PING model of Börgers et al. (2008), gamma results from interactions between excitatory and fast-spiking interneurons. However, tonic inhibition of excitatory and fast-spiking interneurons, mediated by another interneuron population, can facilitate gamma by preventing high firing rates (Börgers et al., 2008). Several findings on visually-induced gamma are compatible with this model: (1) Sst interneurons indeed inhibit both excitatory neurons and PV interneurons (Pfeffer et al., 2013). (2) As shown here, they exhibit relatively weak gamma phase-locking. (3) Gamma amplitude increases with stimulus size and stimulus predictability (Vinck and Bosman, 2016; Uran et al., 2022; Veit et al., 2017). These factors also increase Sst-interneuron firing rates, but decrease firing in PV and excitatory neurons (Adesnik et al., 2012; Keller et al., 2020). Furthermore, suppression of Sst interneurons reduces gamma amplitude (Veit et al., 2017).

In the Borgers/Kopell model, PV interneurons act as the temporal pacemakers of gamma that control the timing of spikes in excitatory cells, and ensure E/I balance and network stability (Okun and Lampl, 2008; Perrenoud et al., 2016). Sst interneurons, on the other hand, would control the amplitude of gamma oscillations by controlling the excitability of PV interneurons and excitatory cells. Such mechanism may enhance the microcircuit’s capacity for the control of gamma oscillations via topdown feedback and behavioral state (Chen et al., 2015; Batista-Brito et al., 2018; Gentet et al., 2012; Veit et al., 2023; Millman et al., 2020).

(2) *Double PING model*. The observation that Sst interneurons fire at a specific phase of the gamma cycle, with a substantial delay after PV interneurons, suggests that they may play a more specific role in providing timed dendritic inhibition. Delayed firing of Sst interneurons compared to PV interneurons may be a more general phenomenon that holds true w.r.t. stimulus onsets (Yu et al., 2019; Estebanez et al., 2017; El-Boustani and Sur, 2014) (see Fig. 1). We further observed that Sst interneurons tended to fire after the occurrence of burst spikes in excitatory neurons. This may suggest that Sst interneurons are particularly activated by these burst spikes, due to the facilitatory nature of E-to-Sst synapses. In turn, studies suggest that activation of Sst interneurons suppresses bursting in excitatory neurons, by inactivating the dendritic compartments involved in burst-firing (Gentet et al., 2012; Royer et al., 2012; Murayama et al., 2009).

The relation between Sst interneurons and burst-firing in excitatory cells suggests a “double PING” model: In the gamma cycle, the initial activation of excitatory neurons leads to rapid recruitment of PV interneurons, which leads to a subsequent decrease in the firing of excitatory neurons through strong perisomatic inhibition. However, the initial activation of excitatory neurons causes back-propagating (sodium) action potentials that further depolarize the dendrites and can cause a burst of action potentials, possibly mediated by voltage-gated active conductances (Larkum et al., 1999; Wang, 1999). Around this time in the gamma cycle, the delayed activation of Sst interneurons increases dendritic inhibition, which suppresses burst firing. In this “Double PING” model, PV and Sst interneurons play complementary roles in controlling the excitability of somatic and dendritic neural compartments, respectively, with precise time delays. Sst interneurons may contribute to the stabilization of gamma-rhythmic dynamics by the timed inhibition of dendritic compartments and control of burst firing (Gentet et al., 2012).

There is increased evidence of a strong association between visually induced V1 gamma oscillations and burst firing: We found burst spikes are more phase-locked than single spikes in mouse V1, and macaque V1 contains a specialized cell type of bursting excitatory neurons with very strong gamma phaselocking (Onorato et al., 2020; Gray and McCormick, 1996). The control of burst firing may be functionally significant for information processing and learning. Bursting may be a specialized cellular mechanism to ensure reliable transmission of information across areas (Lisman, 1997), but is also implied in learning (Lisman, 1997; Doron et al., 2020).

### Subclasses of Sst interneurons

We identified two subclasses of Sst interneurons:

(1) A subclass of Sst interneurons with narrow waveforms and similar phase-locking strength as PV neurons, but with delayed firing in the gamma cycle. Hence, these Nw-Sst neurons make a relatively strong contribution to the rhythmic inhibition of pyramidal neurons’ dendrites. This sub-class of Sst neurons may overlap at least partially with the Non-Martinotti cells. Indeed, NMC tends to have a narrow waveform shape and represents roughly the proportion of Nw Sst we found in our dataset (28%) (Nigro et al., 2018; Muñoz et al., 2017; Kim et al., 2016; Kvitsiani et al., 2013). Although Sst-Nw neurons had strong gamma phase-locking, they fired at a late phase in the gamma cycle, which is likely explained by their facilitating properties (Naka et al., 2019).

(2) The larger class of Bw Sst interneurons, should mostly overlap with the Martinotti type. Those Marinotti cells are the main Sst group (Rudy et al., 2011b; Muñoz et al., 2014; Nigro et al., 2018) and present long axonal plexus that target distal dendrites of pyramidal cells in the superficial layers (Dipoppa et al., 2018; Pfeffer et al., 2013; Muñoz et al., 2017). The relatively weak gamma phase-locking in Bw Sst interneurons agrees with a recent study that found a lack of phase-locking to 40 Hz visual flicker stimuli in Bw Sst interneurons compared to Nw Sst and PV interneurons (Schneider et al., 2023).

Few histological studies reported in the Sst-Cre line an expression leakage into the PV population (Hu et al., 2013). Yet, the proportion of identified Nw Sst cells in our preparation is higher than the range of reported off-expression from the CRE-lines (2-10%) (Hu et al., 2013). Moreover, the gamma-phase delay between Nw-Sst and PV neurons suggests that these are different cell classes.

### Outlook

In sum, our data suggests that the synchronization of excitatory neurons mainly results from the rhythmic activity of PV interneurons. Sst interneurons show less temporal precision and fire with a systematic delay relative to PV interneurons, and may therefore provide timed control of dendritic excitation coordinated at the time scale of gamma cycles. This suggests a model of gamma that goes beyond the classic PING model, and incorporates the morphology of excitatory neurons. In this “Double PING” model, perisomatic and dendritic inhibition are coordinated with distinct spiking events in excitatory neurons.

A systematic phase delay between perisomatic and dendritic inhibition could mean that in the gamma-cycle, there are different windows of opportunity for *firing* and for *learning*. The window of firing would follow the rhythmic inhibition mediated by PV interneurons, which broadly gates the outputs of excitatory neurons at the soma. By contrast, the window for learning would be set by the rhythmic modulation of Sst interneurons, which regulate the voltage membrane potential at excitatory synapses in the dendrites that are known to play important roles in learning (Koch, 2004; Adler et al., 2019). The function of gamma-rhythmic dendritic inhibition, mediated by Sst interneurons, may go beyond the gating of spiking output *per se*, as dendritic depolarization by itself is a key factor for synaptic plasticity (Golding et al., 2002; Sjöström and Häusser, 2006).

## Acknowledgements

We thank Dr. Quentin Perrenoud, Dr. Renata Batista-Brito, Dr. Georgios Spyropoulos, and Dr. Andres Canales-Johnson for helpful comments.

## Authorship Contributions

Conceptualization and design: IO, AB, MV. Analysis main data set: IO. Analysis Allen data: MS. Recordings: IO. Surgery: IO, AT. Materials and reagents: AT, CU, MV. Supervision: MV. Funding acquisition: MV. Writing initial draft: IO, MV. Review/Editing: AB, AT, MS, CU.

## Methods

### Transgenic mice

Experiments were performed on three to eight months old mice, both genders were used. All procedures complied with the European Communities Council Directive 2010/63/EC and the German Law for Protection of Animals and were approved by local authorities, following appropriate ethics review. Mice were socially housed with their litter on an inverted 12/12 h light cycle and recordings were performed during their dark (awake) cycle. To identify the PV-positive (PV) and Sst-positive neurons (Sst) during electrophysiological recordings we crossed the genetic mouse line Ai32(RCL-ChR2(H134R)/EYFP), which contains a CRE-dependent ChannelRhodopsin-2 (ChR2), with either PV-Cre mice (B6.129P2-PVtm1(cre)Arbr/J, JAX Stock 017320, The Jackson Labaratory) or Sst-IRESCre mice (Ssttm2.1(cre)Zjh, JAX Stock 013044, The Jackson Laboratory), to allow Cre-dependent expression of ChR2 in PV (PV-ChR2) and Sst neurons (Sst-ChR2), respectively.

### Preparation for in vivo recording

Thirty minutes prior to the head-post surgery antibiotic (Enrofloxacin, 10 mg/kg, sc, Bayer, Leverkusen, Germany) and analgesic (Metamizole, 200 mg/kg, sc) were administered. For the anesthesia induction, mice were placed in an induction chamber and briefly exposed to isoflurane (3% in oxygen, CPPharma, Burgdorf, Germany). Shortly after the anesthesia induction, the mice were fixated in a stereotaxic frame (David Kopf Instruments, Tujunga, California, USA) and the anesthesia was adjusted to 0.8 – 1.5% in oxygen. To prevent corneal damage the eyes were covered with eye ointment (Bepanthen, Bayer, Leverkusen, Germany) during the procedure. A custom-made titanium head fixation bar was secured with dental cement (Super-Bond C & B, Sun Medical, Shiga, Japan) exactly above the bregma suture, while the area of the recording craniotomy (V1, AP: 1.1 mm anterior to the anterior border of the transverse sinus, ML: 2.6 mm) was covered with cyanoacrylate glue (Insta-Cure, Bob Smith Industries Inc, Atascadero, CA USA). Four to six days after the surgery, the animals were habituated for at least five days in the experimental conditions. The day of the first recording session a 0.6 mm^2^ craniotomy was performed above V1 (centered in the V1 coordinates) under isoflurane anesthesia. The craniotomy was covered with silicon (Kwik-Cast, World Precision Instruments, Sarasota, USA), and the mouse was allowed to recover for at least 2 hours. Recording sessions were carried out daily for a maximum of 5 days, depending on the quality of the electrophysiological signal. Awake mice were head-fixed and placed on the radial wheel apparatus. The animals were mostly stationary, with occasional instances of running (12.2% of the total session time).

### Visual stimulation

Visual stimuli were generated using Psychophysics Toolbox (Brainard and Vision, 1997). The experiment was run on a Windows 10 and stimuli were presented on a monitor with a 144 Hz refresh rate. The Square-wave gratings stimulus was presented for 2 seconds interleaved by, on average, 2 seconds of inter-trial interval. The pre-stimulus duration time had a random duration between 1.9 and 2.1 seconds. The stimuli were presented full-screen in 4 randomized directions, full contrast, a spatial frequency of 0.04 cycles per degree, and a temporal frequency of 4 degrees per second. During this time the monitor’s screen color was gray, and was gamma corrected in order to have the same luminance value as the square-wave gratings stimulus.

### Electrophysiological recordings and optogenetics

For the electrophysiological recordings, we used either Neuropixels (animals = 11, opto-tagged cells = 115 cells) or Cambridge Neurotech probes (animals = 9, opto-tagged cells = 44). During the experiments performed with Neuropixels probes, we inserted the probe up to a depth of 1100 μm and recorded simultaneously from 110 channels (Fig. 1a). In the experiments performed with Cambridge Neurotech probes (H2 and H3 models), we recorded from 64 channels simultaneously.

We targeted PV and Sst interneurons using optogenetic stimulation. During the optogenetic experiment, an optic fiber (Thorlabs, 200um, 0.39 NA) coupled to a diode laser (LuxX CW, 473 nm, 100 mW, Omicron-Laserage Laser produkte GmbH, Germany) was used to illuminate V1 craniotomy. The optic fiber was positioned 0.2 mm from the probe position, just above the surface of the brain. Continuous light square pulses were applied for 1 second interleaved by 3-6 s intervals.

## DATA ANALYSIS

### Waveform classification

The mean waveform was calculated over data segments from -41 to 42 samples around the time of the spike, based on the aligned waveforms of the 10000 spikes per unit randomly selected across the whole recording (excluding the laser trials). The sampling rate was increased by a factor of 10 using spline interpolation. The mean waveforms were normalized by subtracting the median of the first 10 samples and then dividing by the absolute value of the negative peak. Waveforms with a positive absolute peak were discarded. Subsequently, two-dimensional t-Stochastic Neighbor Embedding (t-SNE; perplexity of 80) was applied on the samples after the negative spike peak of the waveforms. Lastly, we applied hierarchical clustering on the two-dimensional t-SNE embedding, which resulted in two separate clusters corresponding to the broad and narrow waveform neurons (Fig. 1e). The proportion of broad and narrow waveforms was ≈85 and 15 percent, respectively. Given the different filtering settings used to record with Neuropixels and Cambridge Neurotech probes, the waveforms were clustered separately in the two datasets.

### Assignment of cortical layers in V1

The cortical depth of each channel was assigned by zeroing the channel at the surface of the brain. The laminar probe was inserted into the brain at an angle of 15 degrees. The depth of the recorded units was corrected for the insertion angle accordingly. Each neuron was assigned a depth corresponding to the estimated depth of the channel in which its waveforms presented the highest amplitude.

### Optotagging protocol

Optogenetic tagging experiments were performed on the mice line where an opsin was selectively expressed either in PV or Sst interneurons. The opto-tagging protocol consisted of 40 trials of 1s-long stimulation periods with a randomized inter-stimulus interval between 4 and 6 seconds. Neurons were considered light-responsive if were fulfilling two criteria: (1) A significant increase of firing probability (when the average of spike count is higher than the random permuted matrix) after light onset (excluding the first 0.5ms to avoid the presence of light artifacts). (2) A positive firing rate modulation compared to baseline during the first 200ms of a continuous square light pulse (Fig. 1b-c). The firing modulation was calculated with the following formula: FR(laser)FR(baseline)/FR(laser)+FR(baseline). To test the validity of the opto-tag criteria, we also included an additional criterium to identify light-responsive cells, namely a significant increase of firing probability within the first 5 milliseconds after the light onset. The use of this criteria yielded a smaller number of optotagged interneurons (PV n = 49, Sst n = 41), but resulted in similar conclusions as in the main Figures (Figure S2d-f).

### LFP preprocessing and Spectral analysis

Single units were isolated using the semi-automated spike sorting algorithm Kilosort 2 (Steinmetz et al., 2021). LFP signals were referenced to a channel near the dura, low-pass filtered at 400 Hz, and high-pass filtered at 0.1 Hz, using a third-order Butterworth filter. In order to filter out line noise, an additional band-stop filter between 49.5 and 50.5 Hz was applied. Subsequently, signals were downsampled to 1200 Hz. To estimate the LFP power spectra in the stimulus and baseline periods (Fig. 1i) and supplementary figure S1, d), we used the following procedure: Power spectra were estimated separately for the prestimulus and the visual stimulation period. The duration of the visual stimulus was 2 seconds, for the frequency analysis, we excluded the first and last 200 milliseconds of the stimulus period to remove the neuronal evoked response period and possible anticipatory effects, respectively. Data epochs were Hanning tapered, to achieve a fundamental spectral resolution (Rayleigh frequency) of 2 Hz and then Fourier transformed.

To compute spike-LFP phase-locking, we determined the phase of each spike relative to the LFP channel presenting the highest power at the gamma frequency range (25-40 Hz), making sure it has a distance of at least 400 um from the spike channel.

To do so, we computed the wavelet transform of the LFP snippet around each spike, using a constant number of cycles (9) per frequency (as in (Vinck et al., 2013)), using the “ft spiketriggeredspectrum” functions in the FieldTrip SPIKE toolbox and after computing the spike-LFP phases, we estimated phase-locking with the pairwise-phase-consistency (PPC) (Vinck et al., 2010, 2012). Specifically, we used the PPC1 measure defined by (Vinck et al., 2012). The PPC1 takes all pairs of spikes-LFP phases from separate trials and computes the average consistency of phases across these pairs. It is not affected by mean discharge-rates and history effects like bursting (Vinck et al., 2012) (Fig. 2c and Fig. 4b and Fig. 3c). For more details see (Onorato et al., 2020). Because phase-locking estimates can have a high variance for low spike counts, we computed PPC values only for neurons that fired at least 100 spikes (Vinck et al., 2013) (Fig. 2d and Fig. 4c and Fig. 3a). To estimate the spikes preferred phase we only included neurons that presented an average value of phase-locking at the gamma frequency-range (25-40Hz) higher than 0.002.

We occasionally observed the emergence in V1 of stereotypical events at 4-8 Hz. Studies have reported that this rhythm may affect the visual processing in V1 (Gao et al., 2021; Speed et al., 2019). Hence, we excluded, for the spectral analysis, the trials with high power in the 4-8 Hz frequency range of the Fourier transform.

### Quantification of spiking data

To estimate the neuronal visual response we computed the normalized peri-stimulus-time-histogram (PSTH, binning = 0.005 s) diving the average trial response by the mean baseline firing rate (Fig. 1d). The delay of the neuronal response to the stimulus onset was found as the time latency of the normalized PSTH to reach a threshold 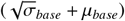 (Fig. 1f).

Spikes were labeled as burst spikes if had an interval to the next spike longer than 6 milliseconds (Fig. 3c, top plot), while non-burst spikes had to present an interval to the next and the previous spike longer than 10 milliseconds. Non-burst spikes were selected only from the subset of neurons that presented burst events so that burst and non-burst spikes were sampled from the same group of neurons (Fig. 3c, top plot).

The correlation between the neuronal firing rate and the power at different LFP frequencies was calculated as follows: The average firing rate and LFP power were calculated iteratively on a sliding window of 0.4 seconds (from 0.2 to 1.8 seconds after stimulus onset), the LFP power was estimated with the envelope of the Hilbert transform measured on the bandpass filtered LFP. We then computed the Spearman’s rank correlation coefficient across trials independently for each time interval and then we averaged them. We repeated this analysis from 5 to 90 Hz with intervals of 5 Hz (Figures 2e, 4d).

In addition, we computed the Spearman’s rank correlation coefficient between the number of spikes occurring in the interval between the trough and peak of a given gamma cycle and the time interval to reach the peak of the next gamma cycle. The instantaneous gamma phase was detected by computing the four-quadrant inverse tangent of the Hilbert transform of the LFP signal during the stimulus period. This returns a phase signal, whereas the minimum values correspond to the through of each cycle, the peak was identified as the maximum value in the raw LFP between two throughs.

---

**Figure S1:**
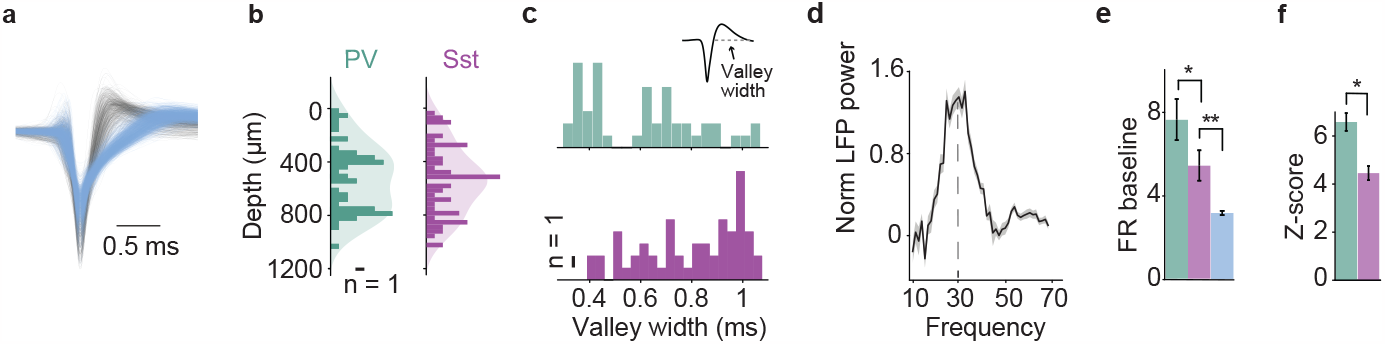
Further analyses related to Figure 1, main text. **a** Action potential waveforms were divided into broad(Bw, n = 2156, light blue) and narrowwaveform (gray, n = 397) groups. For visualization only a subset of cells are shown. **b** Laminar depth distribution of PV (green) and Sst (magenta) interneurons. **c** Distribution of the waveforms’ valleys width. PV interneurons have a shorter valley width than Sst interneurons (p = 0.0029, two-sample t-test). **d** LFP power in stimulus period divided by LFP power in baseline power, shown at a log-scale (20 mice). **e** Average baseline firing rates are higher for PV than Sst (p = 0.04), and higher for Sst than Bw (p = 1.61 ×10^−6^; two-sample t-test). **f** Shown are the Z-scores (max-μ/σ) of the normalized (divided by the baseline) peri-stimulus-timehistogram (PSTH) response to the visual stimulus. Z-scores were computed over the entire trial period, i.e. -2 to +2 s after visual stimulus onset. PV cells are more visually responsive than Sst interneurons (p = 7.1 × 10^−6^; two-sample t-test).

**Figure S2:**
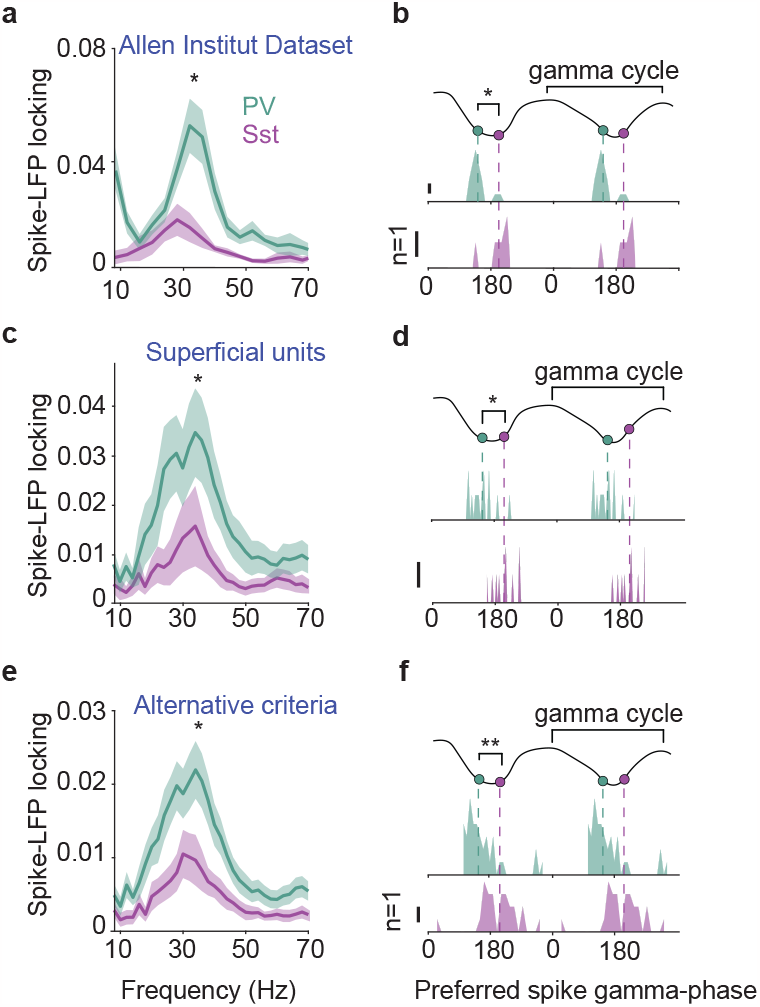
PV has stronger gamma-phase-locking then Sst interneurons. **a** Allen Brain Observatory dataset (Siegle et al., 2021). Average spikes-LFP phase locking (PPC) for opto-tagged PV (n = 12, green) and Sst (n = 5, magenta) interneurons. PV interneurons showed a stronger phase locking in the gamma frequency range (difference at 25-30 Hz: p = 0.0437; two-sample t-test). These recordings were made via Neuropixels during drifting-grating stimulation while mice were placed on a running disk. Similar mice lines were used as in our main dataset (PVand SOM-Cre). For a more detailed description of methods see (Spyropoulos et al., 2022). **b** Distribution of preferred gamma phase of PV and Sst interneurons from the same dataset used for **a**. PV interneurons fired significantly earlier in the gamma cycle than Sst interneurons (p <0.01; Permutation test). **c** Average spikes-LFP phase locking (PPC) for PV (n = 18, green) and Sst (n = 21, magenta) interneurons located in superficial layers (depth <400μm). PV interneurons showed a stronger phase locking in the gamma frequency range (at 25-40 Hz, p = 0.0397; two-sample t-test). **d** Distribution of the preferred gamma phase of PV (n = 13) and Sst (n = 10) interneurons selected as in **c**. The spikes of PV (top) interneurons fired significantly earlier in the gamma cycle than Sst (bottom, p <0.005; Permutation test). **e** Average spikes-LFP phase locking (PPC) for PV (n = 49, green) and Sst (n = 41, magenta) interneurons. In this case, alternative selection criteria for opto-tagging were used that yielded a smaller sample. PV interneurons showed stronger gamma-phase locking than Sst interneurons (at 25-40 Hz: p = 0.0085). **f** Distribution of preferred spike gamma phases of PV and Sst interneurons from the same single units dataset as **e**. PV interneurons fired significantly earlier in the gamma cycle than Sst interneurons (p < 0.0005; Permutation test). **a-f** Error bars and shaded lines indicate s.e.m.

**Figure S3:**
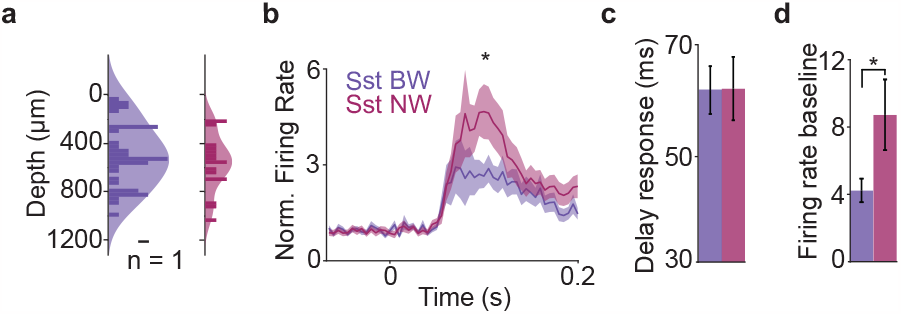
Characterization of Bw and Nw Sst. **a**) Depth distribution of Bw (purple) and Nw (magenta) Sst interneurons. **b**) Average peri-stimulus-time-histogram (PSTH), normalized by baseline firing rate. Nw Sst interneurons has a stronger evoked resopnse than the Bw Sst interneurons (100 ms after stimulus onset, p = 0.02; two-sample t-test, shaded line denotes mean± s.e.m). **c**) Population average of the stimulus latency response. Both Bw and Nw Sst interneurons increased their firing response on average at 60 ms after the stimulus onset (Bw vs. Nw: p = 0.9; two-sample t-test). **d**) Average firing rates in the baseline interval between -1.8 and -0.2 seconds before stimulus onset. Nw has higher baseline firing rates than Bw Sst (p = 0.009; two-sample t-test). **a-d** Error bars and shaded lines indicate s.e.m.

